# Structural presentation of amyloid β (Aβ) by HLA

**DOI:** 10.64898/2026.07.07.736863

**Authors:** Elena Erausquin, María Gilda Dichiara-Rodríguez, Leire Oyon-Olea, Jacinto López-Sagaseta

**Affiliations:** Unit of Protein Crystallography and Structural Immunology, Navarrabiomed, Navarra, Spain; Public University of Navarra (UPNA), Pamplona, Navarra, Spain; Navarra Hospital Complex (CHN), Pamplona, Navarra, Spain

## Abstract

HLA-DR-restricted T-cell reactivity to amyloid β (Aβ) has been associated with Alzheimer’s disease (AD), but structural evidence for HLA presentation of Aβ-derived peptides remains elusive. We present the crystal structure of the Aβ1-15 fragment bound to HLA-DR1, providing, to the best of our knowledge, the first experimental structure of an Alzheimer’s Aβ peptide bound to an HLA molecule. The molecular architecture of this complex defines a peptide:MHC interaction dictated by engagement of Aβ1-15 peptide central core with further involvement of N- and C-terminal peptide flanks. The structure reveals that DRβ1 Arg70, a polymorphic position, directly binds P4 and P5 through polar contacts, providing a rationale for HLA-DRB allelic bias underpinning accommodation of Aβ1-15. We also describe the Aβ1-15:HLA-DRB1 surface topology, informing a candidate binding surface for potential T-cell recognition. Collectively, these findings contribute a structural framework for further research in the context of Aβ-specific CD4^+^ T-cell autoreactivity in AD.

## Introduction

Deposition of amyloid-β (Aβ) peptide, along with tau pathology, neuroinflammation and progressive cognitive decline constitute the hallmark of Alzheimer’s disease (AD)^1^. Although originally AD was understood in the context of neuron disorder and aggregation of local protein, there is mounting evidence that characterize AD by a complex interplay involving resident brain cells and the peripheral immune system. GWAS, single-cell transcriptomics and other experimental approaches associate the immune system with the pathophysiology of AD^2,3^. This includes the microglia, infiltrating immune cells and neuroinflammation mediated by cytokines.

An approach based on transcriptomics provided this profile by unveiling the immune signatures linked to AD at a resolution of single cell^3^. The authors found that in the human prefrontal cortex, AD pathology correlates with diverse subsets of microglial populations and gene modules enriched for immune factors. One such factor is the HLA class II. More specifically, microglial HLA-DRB1, HLA-DRB5 and CD74 genes were found to associate with signs of AD pathology, suggesting that antigen presentation represents a sustained component of the myeloid response in AD^3^. However, the nature of the antigens, the restriction enforced by HLA and the specificity of the T cell response remain to be fully elucidated.

Aβ represents a plausible antigen candidate in this context. Human and mice T-cell reactivity to Aβ have been reported, with DR15, DR3, DR4, DR1 and DR13 being the most prevalent alleles in a cohort of 133 individuals, of which 81 were AD patients^4^. In support of the hypothesis that Aβ-derived peptides can be presented to T cells in an HLA-II-restricted manner, the authors found that T-cell reactivity was directed primarily, but not exclusively, to the Aβ15-42 region^4,5^.

Additional studies by Kutzler and colleagues strengthened the concept of Aβ T-cell recognition mediated by HLA-DR1^6^. Immunization of mice humanized to express HLA-DRB1*01:01 with DNA encoding for human Aβ42 led to IFN-γ production, with the immunodominant CD4^+^ T cell epitope located within Aβ residues 1-28. Therefore, while studies with human patients underscored the central and C-terminal Aβ regions as the triggers for T-cell response, in vivo studies with transgenic mice expressing the HLA-DRB1*01:01 allele reveal that the Aβ N-terminal segment can also elicit an HLA-II-restricted adaptive immune response.

Yet, and regardless of these findings, the current scenario does not provide an experimental structure that confirms and defines a coherent architecture of an Aβ fragment bound to an HLA molecule. Which Aβ residues are configured as MHC anchor points and which residues configure a potential TCR-accessible pMHC topology are questions that remain elusive.

To address this question, we interrogated whether any of the previously identified HLA-DR alleles^4^ are good predictive candidates for presentation of Aβ peptides. NetMHCIIpan 4.2 predicted 15-mer peptide binders with different degree of affinity for all the alleles except DRB1*13:02. Importantly, Aβ1-15 was delivered as the strongest binder with FRHDSGYEV as the 9-mer core motif. Similarly, DRB1*03:01 and DRB1*04:01 were also predicted to accommodate the Aβ1-15 peptide. In contrast, HLA-DRB1*15:01 was predicted to allow binding of the Aβ(17-25) LVFFAEDVG core motif. Therefore, the in silico predictive approach suggested that the Aβ1-15 motif represented a reasonable Aβ candidate for HLA-DR1 presentation, with HLA-DRB1*01:01 showing the strongest affinity and constituting a rationale for the X-ray structural approach.

Here, we present the structure of HLA-DRB1 with bound Aβ 1-15 at a resolution of 2.0 Å, and provide the peptide binding register, the HLA:peptide interaction map and surface topology as a candidate for T-cell recognition.

To the best of our knowledge, this structure represents the first experimental structure of an Aβ-derived fragment presented by an HLA class II molecule and provide a structural rationale for Aβ recognition by CD4^+^ T-cells in an HLA-restricted manner.

## Results

### Aβ1-15 is the strongest predicted HLA-DR1 binder

To identify the most plausible Aβ fragment candidates for structural studies, we used NetMHCIIpan 4.2 as a predictor for the binding of 15-mer Aβ-derived peptides. Aβ(1-42) was the source of peptides, whereas several HLA-DRB1 alleles were considered given their association with Aβ-mediated T-cell reactivity or microglial HLA correlation with AD pathology (Table 1). A series of binders were predicted for most of the HLA-DRB1 alleles, except for HLA-DR1*13:02. Binder potency was dependent on allelic determinants. HLA-DRB1*15:01 yield binders belonging to the Aβ-central segment, with common core settled in the LVFFAEDVG motif. On the contrary, HLA-DRB1*03:01 and HLA-DRB1*04:01 Aβ predicted binders converged in the N-terminal Aβ1-15 region, which is defined by the DAEFRHDSGYEVHHQ sequence and the 9-mer core FRHDSGYEV. With a predicted binding affinity of 64 nM, the binder showing the strongest interaction was found for HLA-DRB1*01:01, again, with Aβ target sequence Aβ1-15. An analogous peptide, shifted to yield Aβ(2-16), was also estimated to bind HLA-DRB1*01:01 using the same core and a predicted affinity of 58 nM. The %Rank_EL, however, delivered values of 0.49 and 1.60 for Aβ1-15 and Aβ(2-16), respectively. Because strong candidates show rank values below 0.5%, Aβ1-15 was chosen as the best binder candidate for X-ray studies.

**Table 1.**
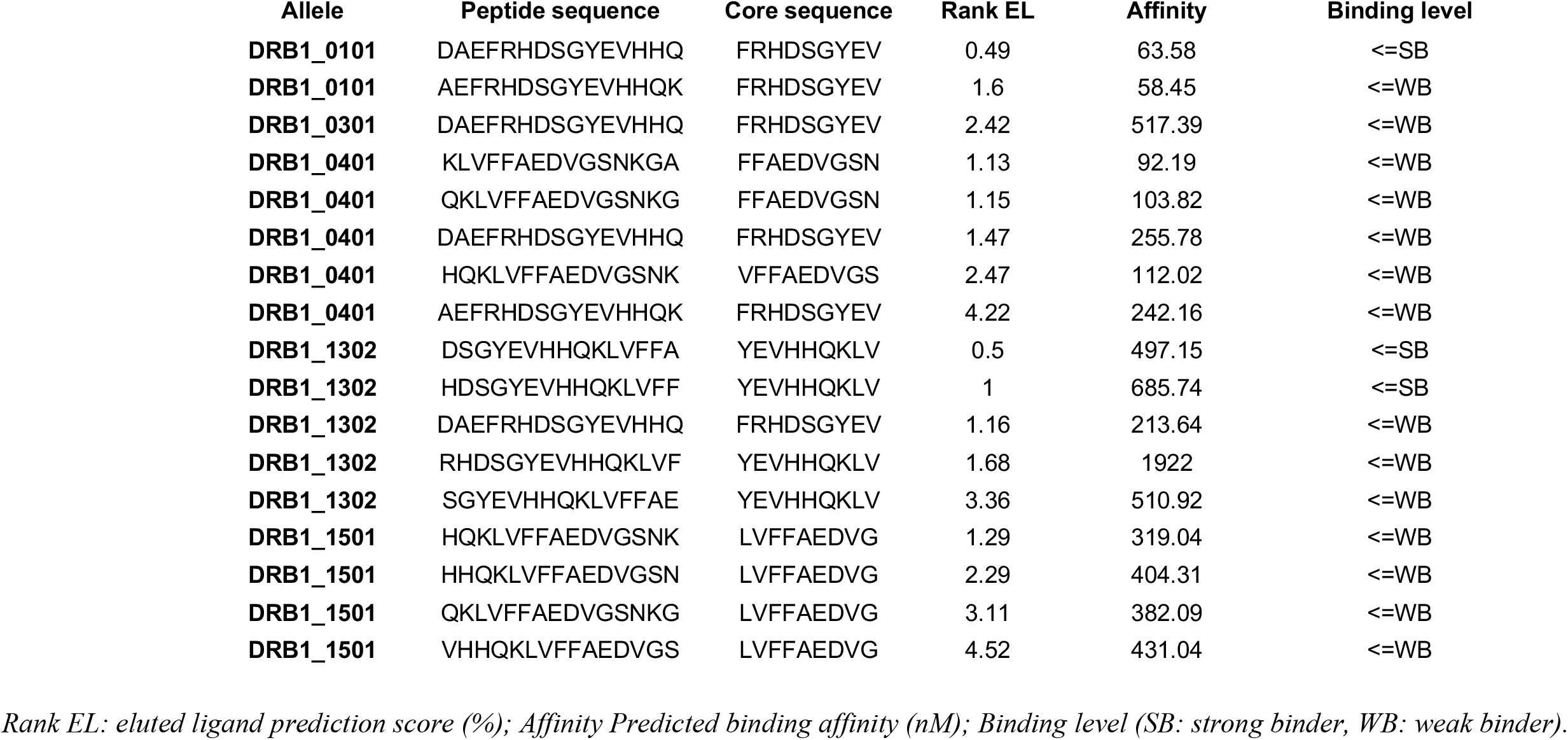
Prediction of binding of Aβ42-derived peptides to selected HLA-DRB1 alleles.

### Crystal structure of HLA-DR1 with bound Aβ peptide 1-15 fragment

We solved the crystal structure of HLA-DRA/HLA-DRB1*01:01 with bound N-terminal Aβ peptide 1-15 fragment. The rationale behind this aim was founded on prior findings linking AD pathology to microglial HLA-DR1. The lack of experimentally determined structural evidence for Aβ presentation by HLA further inspired this work (Fig. 1a and Table 2).

**Table 2.** Diffraction data collection and refinement statistics.

| **DATA COLLECTION** | **Aβ1-15:HLA-DR1\*01:01** |
| --- | --- |
| **Resolution range (Å)** | 55.91 - 2.00 (2.05 - 2.00) |
| **Space group** | P12<sub>1</sub>1 |
| **Unit cell** | 57.31 118.51 67.13 90.00<br>109.15 90.00 |
| **Total reflections** | 387514 (23599) |
| **Unique reflections** | 56684 (4148) |
| **Multiplicity** | 6.8 (5.7) |
| **Completeness (%)** | 99.3 (98.2) |
| **Mean I/sigma(I)** | 7.7 (1.9) |
| **Wilson B-factor** | 32.63 |
| **R-merge** | 0.153 (1.580) |
| **R-meas** | 0.165 (1.75) |
| **R-pim** | 0.063 (0.737) |
| **CC1/2** | 0.993 (0.395) |
| **Reflections in refinement** | 56550 (2673) |
| **Reflections (R-free)** | 2902 (139) |
| **R-work** | 0.1849 (0.239) |
| **R-free** | 0.2203 (0.306) |
| **RMS(bonds)** | 0.010 |
| **RMS(angles)** | 0.92 |
| **Ramachandran favored (%)** | 99.19 |
| **Ramachandran allowed (%)** | 0.81 |
| **Ramachandran outliers (%)** | 0.00 |
| **Average B-factor (Å<sup>2</sup>)** | 40.28 |

Statistics for the highest-resolution shell are shown in parentheses.

**Figure 1.**
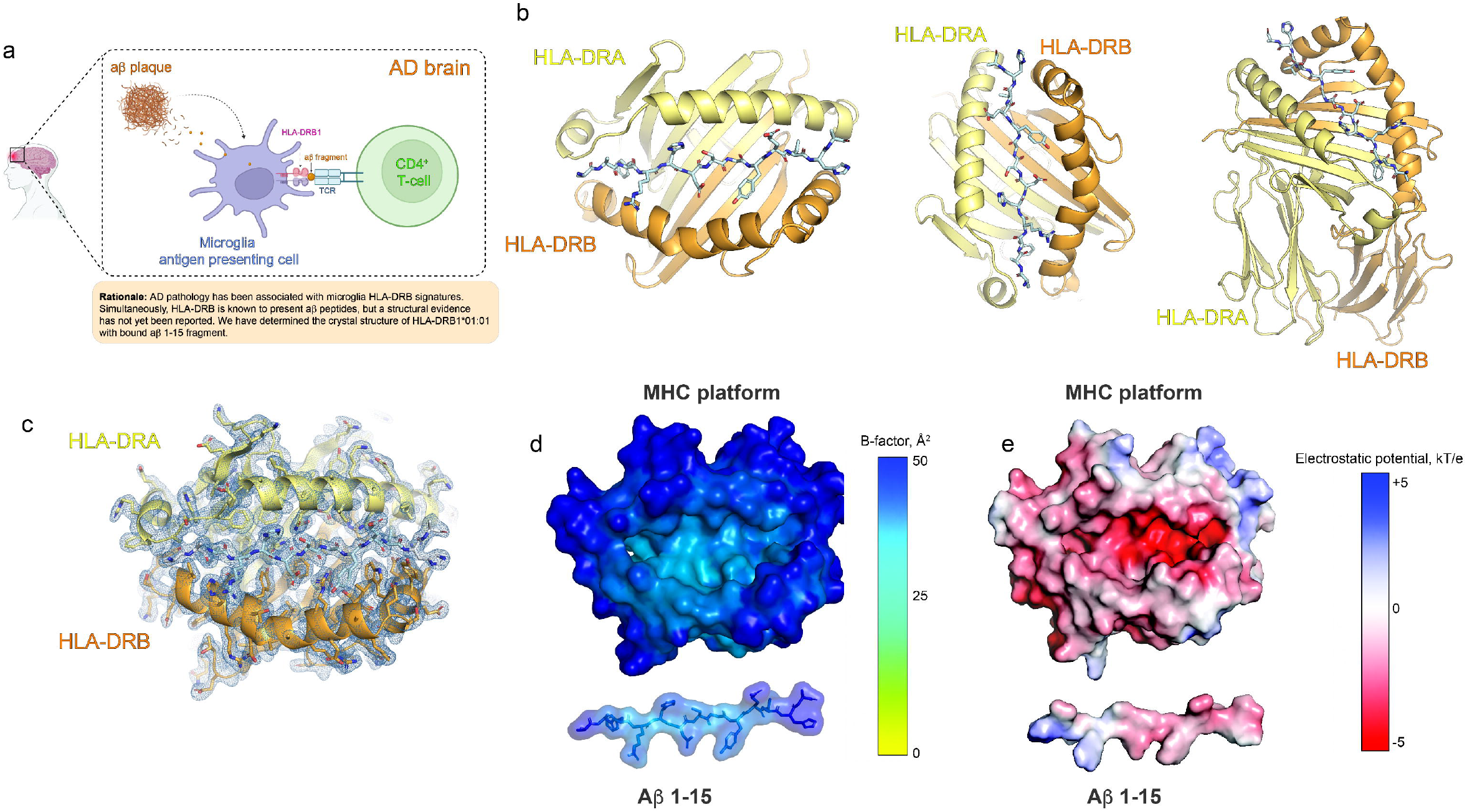
Aβ1-15:HLA-DR1 crystal structure. **A** Rationale underpinning the assessment of HLA-DR1 structural competence to bind an amyloid-β-derived peptide. A positive correlation has been described for microglial expression of HLA-DR1 and Alzheimer’s disease pathology. This raised the question of whether Aβ could lead to peptide fragments structurally suitable for presentation to CD4^+^ T-cells by HLA-DR. **B** Three alternative views of the Aβ1-15:HLA-DR1 crystal structure complex. HLA-DRA is shown in pale yellow, HLA-DRB1 in orange and the Aβ1-15 peptide highlighted as sticks. **C** Composite 2Fo-Fc electron-density map (1 sigma contour level) of one of two Aβ1-15:HLA-DR1 complexes present in the assymmetric unit. **D** B-factor scan for the Aβ1-15:HLA-DR1 complex. Upper panel, MHC platform. Lower panel, Aβ1-15. **E** Electrostatic surface potential for the Aβ1-15:HLA-DR1 complex. Upper panel, MHC platform. Lower panel, Aβ1-15.

The HLA-DR1:Aβ1-15 presents a classical pMHC(II) architecture, with the peptide anchored in the central cleft defined by HLA-DRA and HLA-DRB1 chains (Fig. 1b). We obtained crystals in P1211 space group, with the asymmetric unit presenting two HLA-DRB1:Aβ 1-15 complexes. Data processing and refinement was accomplished at 2.0 Å resolution with Rwork and Rfree values of 0.189 and 0.220, respectively (Table 1).

Fo-Fc and 2Fo-Fc electron density maps guided building of the peptide and the MHC platform, with 2Fo-Fc showing continuous density for the alpha and beta chains as well as for the bound peptide (Fig. 1c). The MHC and peptide molecules present high internal order according to the B-factor analysis, and together with the electron density signal enabled precise location of the peptide register and trajectory (Fig. 1d).

An electrostatic view of the complex indicates that Aβ1-15 is bound in a negatively charged HLA cleft, with Aβ1-15 displaying a heterogeneous electrostatic topology characterize by a short positively charged N-terminal region and a longer, negatively charged C-terminus (Fig. 1e).

These data settle a precise atomic view of the HLA-DR1:Aβ1-15 complex, providing the first experimentally determined molecular assembly of an Aβ fragment being presented by an HLA class II molecule.

### Aβ1-15 is found within HLA-DR1 groove with a defined peptide binding register

The Fo-Fc signal allowed unbiased assignment of the Aβ1-15 residues within HLA-DR1 groove (Fig. 2). The shape of the residues in the RHDSGY motif of Aβ1-15 were particularly supportive to identify the correct peptide binding register (Fig. 2a). The resulting 2Fo-Fc density map was coherent with the inferred peptide register and orientation of the side chains. The precise arrangement of the Aβ1-15 fragment followed a register with the N-terminal DAE residues (<u>DAE</u>FRHDSGYEVHHQ) creating the N-flanking site, with poor density signals for the side chains for AE, or absent in the case of the first position, Asp (<u>D</u>AEFRHDSGYEVHHQ). Following this segment, Aβ1-15 locates the 9-aa core defined by F4, R5, H6, D7, S8, G9, Y10, E11 and V12 (FRHDSGYEV). This motif corresponds to positions P1 through P9 of the HLA-DR1 groove (Fig. 3a).

**Figure 2.**
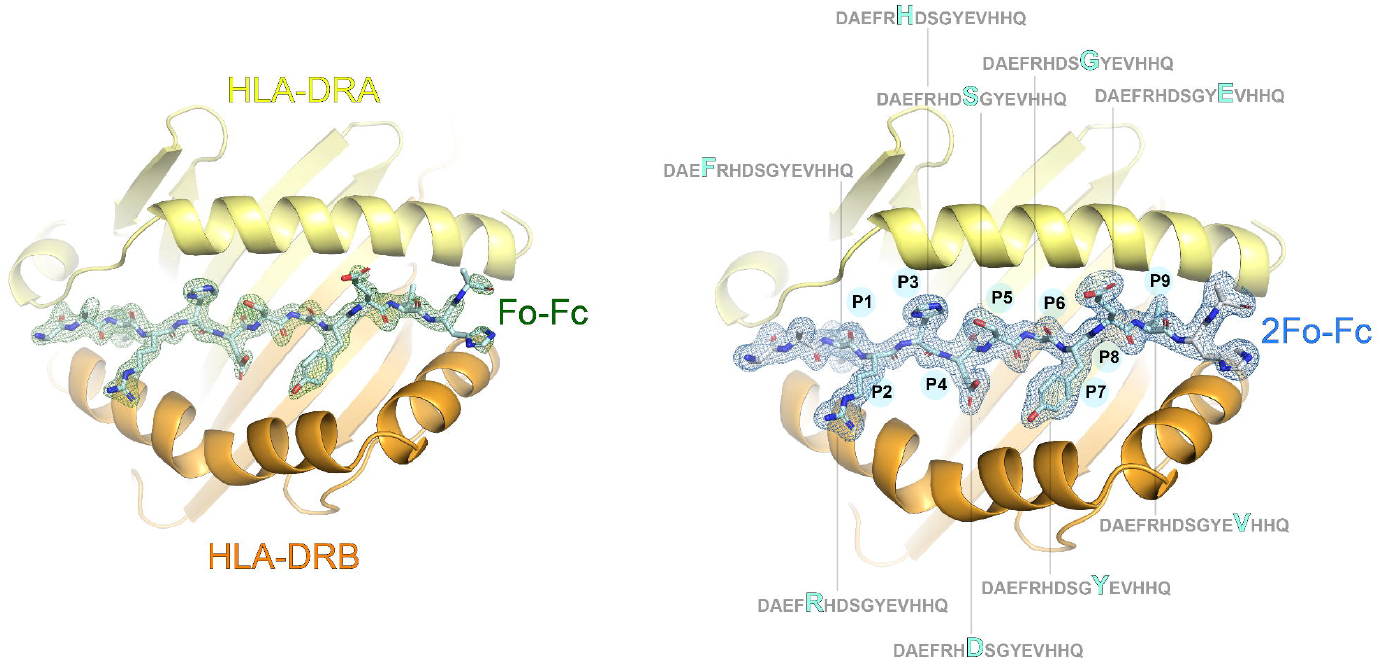
Electron density maps and peptide binding register of Aβ1-15. **A** Omit Fo-Fc electron-density map (contoured at 3 sigma) observed within HLA-DR1’s groove. **B** One sigma-contoured 2Fo-Fc electron-density map obtained upon refinement for Aβ1-15 bound to HLA-DR1. Aβ1-15 residues forming the canonical 9-mer core are labeled.

**Figure 3.**
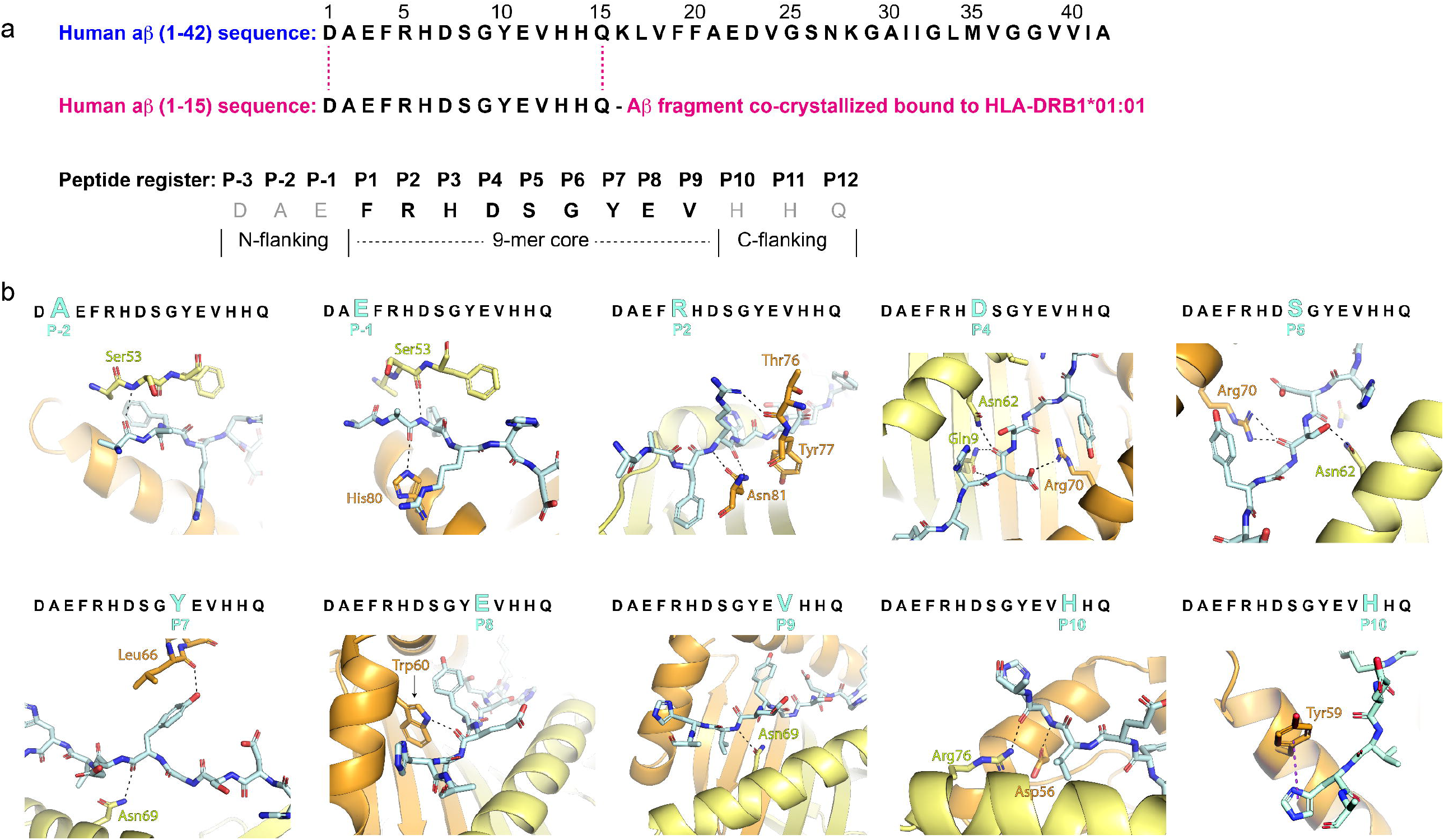
Aβ1-15 peptide binding register within HLA-DR1 and molecular engagement. **A** Sequence of Aβ42 and that of the crystallized Aβ1-15 fragment. The peptide-binding register is displayed and split into core and flanking regions. **B** Close-up views of the observed polar Aβ1-15:HLA-DR1B contacts (dashed lines), distributed along the MHC’s groove.

The Aβ1-15 trace within HLA-DR1 ends with the C-terminal flanking region formed by residues HHQ. As for the N-terminal segment, we observe weak signal for second histidine side chain but complete absence of signal for the Gln residue.

Altogether, these observations support a tightly bound central Aβ1-15 core defined by a 9-aa motif, and flexible flanking regions at both N- and C-termini, which is consistent with an open-ended HLADR1 architecture of its peptide binding groove.

### HLA-DR1:Aβ1-15 is stabilized through primary polar contacts across the peptide core and auxiliary flanking residues

We examined the HLA-DR1:Aβ1-15 binding signature. Aβ1-15 forms a dense network of polar interactions with residues located in both HLA-DRA and HLA-DRB1 (Fig. 3b and Supplementary Table 1), including contacts mediated by residues beyond the central P1-P9 core. More specifically, the N-flank establishes backbone interactions with HLA-DR1 that involve peptide residues Glu in P-1 and Ala in P-2 (Fig. 3b). As for the C-flank, we observe interactions with P10 (His). These appear to contribute significantly to the overall HLA-DR1:Aβ1-15 assembly, as they involve, one the one hand, contacts with HLA-DRA (Arg76) and HLA-DRB (Asp56) helices through side chains (HLA) and backbone (Aβ fragment) atoms (Fig. 3b), and on the other hand, a π-π interaction formed between the His imidazole ring and HLA-DRB1 Tyr59. This network of interactions established with the HLA molecule suggests a relevant role for the Aβ His residue located in P10 position in its mode of engagement within HLA-DR1 groove.

Within the P1-P9 core, several residues are contacted by HLA-DRA and HLA-DRB chains. Considering polar interactions, Aβ1-15 Arg (P2)’s side chain interacts with DRB Tyr77 backbone, whereas its backbone binds DRB Asn81 side chain. Aβ1-15 Asp (P4) establishes a salt bridge with DRB Arg70 and is further locked into P4 position by backbone polar contacts with DRA Gln9 and Asn62 (Fig. 3b). Ser (P5) interacts also with DRB Arg70 and DRA Asn62. Both HLA chains contact also Aβ1-15 Tyr (P7) through DRA Asn69 and DRB Leu66. Lastly, Glu (P8) and Val (P9) appear bound to DRB Trp60 and DRA Asn69, respectively, via side chain (HLA)-backbone (Aβ) interactions.

### HLA-DRB1 polymorphisms map to residues involved in Aβ1-15 binding

Next, we examined HLA-DRB polymorphisms at the peptide:MHC interface. Alignment of HLA-DRB alleles associated with risk or protection in AD revealed several polymorphic sites located at the HLA peptide binding cleft (Table 3). Position 70 in HLA-DRB is occupied by Arg in HLA-DRB1*01:01 and, as described earlier, is found to participate in multiple contacts with Aβ1-15 Asp in P4 and Ser in P5 (Figs. 3b and 4). In DRB1*13:02 and DRB1*15:01, however, position 70 is occupied with Glu and Ala, respectively, which contribute opposite electrostatics and local geometry. Thus, position 70 occupies a structurally relevant site at the peptide-MHC interface that can influence Aβ1-15 accommodation within HLA-DRB.

**Table 3.** HLA-DRβ polymorphism at the peptide:MHC interface.

|  | DRβ position |  |  |  |  |  |  |  |  |  |  |  |  |  |  |  |  |
| --- | --- | --- | --- | --- | --- | --- | --- | --- | --- | --- | --- | --- | --- | --- | --- | --- | --- |
| DRB1 ALLELE | 55 | 56 | 57 | 58 | 59 | 60 | 61 | 62 | 63 | 64 | 65 | 66 | 67 | 68 | 69 | 70 | 71 |
| DRB1*01:01:01:01 | P | D | A | E | Y | W | N | S | Q | K | D | L | L | E | Q | R | R |
| DRB1*04:01:01:01 | P | D | A | E | Y | W | N | S | Q | K | D | L | L | E | Q | K | R |
| DRB1*04:03:01:01 | P | D | A | E | Y | W | N | S | Q | K | D | L | L | E | Q | R | R |
| DRB5*01:01:01:01 | P | D | A | E | Y | W | N | S | Q | K | D | F | L | E | D | R | R |
| DRB1*15:01:01:01 | P | D | A | E | Y | W | N | S | Q | K | D | I | L | E | Q | A | R |
| DRB1*13:02:01:01 | P | D | A | E | Y | W | N | S | Q | K | D | I | L | E | D | E | R |
| DRB1*04:04:01:01 | P | D | A | E | Y | W | N | S | Q | K | D | L | L | E | Q | R | R |
| DRB1*09:01:02:01 | P | V | A | E | S | W | N | S | Q | K | D | F | L | E | R | R | R |

*Violet labeling indicates residues that contact Aβ1-15 through polar interactions and are conserved across HLA-DRβ. Orange color corresponds to substitutions with similar properties. Black shading designates non-conservative substitutions.*

**Figure 4.**
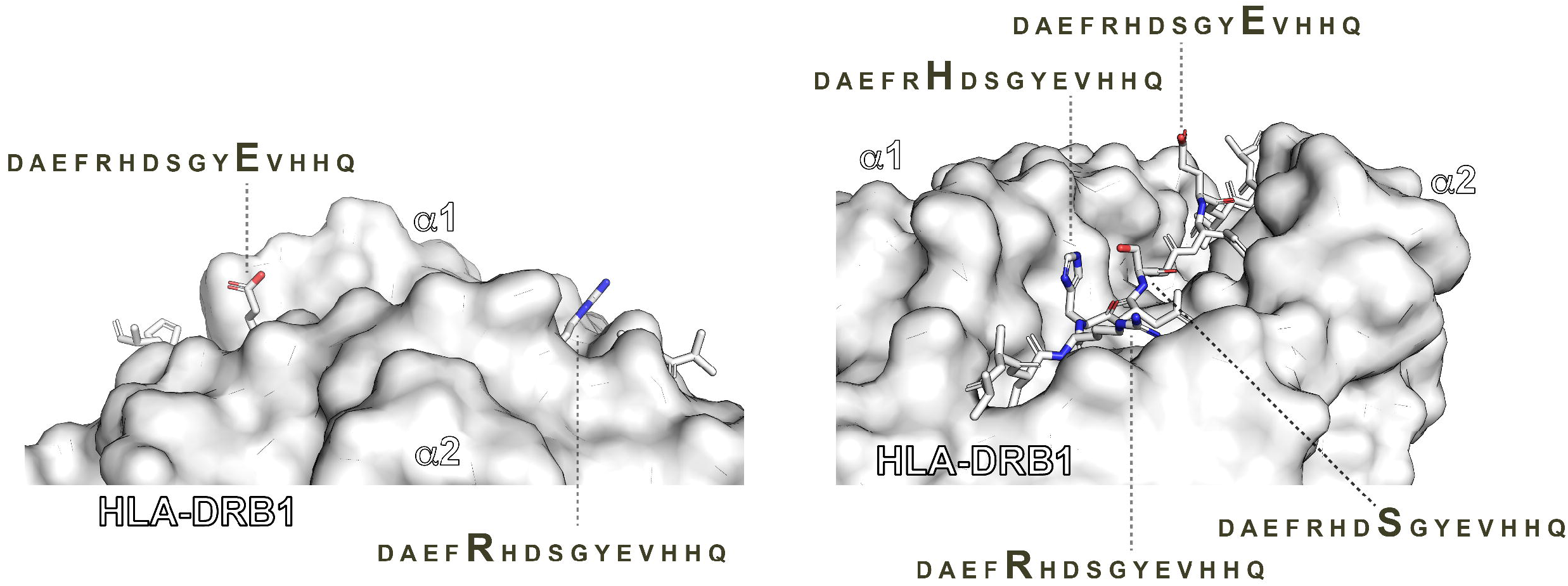
Molecular topology at the Aβ1-15:HLA-DR1 complex surface. **A** Surface representation of HLA-DR1, with the Aβ1-15 peptide highlighted as sticks, either colored or greyed for solvent-exposed and buried residues, respectively. Surface-accesible residues are labeled and highlighted within the Aβ1-15 sequence.

### The Aβ1-15:HLA-DRB1 surface topology defines potential T-cell recognition motifs

To map a putative T-cell recognition motif, we evaluated which residues in Aβ1-15 are solvent-exposed. We found that the side chains of His (P3) and Glu (P8) remain pointing upwards and are not engaged by the MHC (Figs. 3b and 4). Therefore, these residues represent plausible recognition sites for a potential Aβ1-15-reactive T-cell receptor (TCR). Additionally, Arg (P2) and Ser (P5), although involved in intermolecular contacts with the MHC, are solvent-exposed, making them second-line candidates for T-cell engagement.

## Discussion

We report the crystal structure of HLA-DR1 with a bound amyloid-β (Aβ)-derived peptide. The structure describes the binding mode of Aβ1-15 within the peptide binding cleft of HLA-DRB1, with beta chain HLA-DRB1*01:01, reports core and flanking peptide contact residues and determines the peptide register. To the best of our knowledge, this structure represents the first atomic view of an Aβ-derived peptide presented by a human HLA molecule, providing, therefore, a molecular framework for future studies in the context of CD4^+^ T-cell immunity in AD.

Prior studies by several groups support a link between Aβ and HLA class II at a molecular, genetic and immunological level. Studies performed with human samples have informed the presence of T-cell Aβ-reactivity in an HLA-restricted manner and underscoring the immunodominant role of Aβ central- and C-terminal regions^4,5^. Further single-cell transcriptomics of human AD brain specimens provided a positive correlation between the expression of HLA-DRB1 in the microglia and pathologic signs in AD^3^. While these studies did not determine that HLA-DRB naturally presents Aβ-derived peptides in the AD brain, they provide a foundation for investigating the potential for Aβ to become a source of antigens presented by HLA-II.

This hypothesis was supported by computational prediction of Aβ binding to AD-linked HLA-DRB1 alleles. NetMHCIIpan 4.2 found Aβ-derived 15-mer peptide candidates with varied degrees of affinities. Unlike the other alleles, which were predicted to have a preference for the N-terminal Aβ1-15 segment, HLA-DR1*15:01 binders concentrate in the Aβ central region, with the core dominant motif LVFFAEDVG, which corresponds to Aβ(17-25).

Likely the most relevant observation lies in the fact that the stronger binder candidate was found for HLA-DR1*01:01, with a predicted binding affinity and eluted-ligand rank scores surpassing any other HLA-peptide combination. This candidate is defined by Aβ residues 1-15, with the FRHDSGYEV core motif located in the Aβ(4-12) region. Although with poorer scores, this motif was shared with the HLA-DR1*03:01 and HLA-DR1*04:01 alleles. Due to the predicted high affinity and binding probability, Aβ1-15 was the targeted peptide antigen candidate for structural studies along with the HLA-DR1*01:01 allele.

Importantly, previous reports underscored a T-cell response to Aβ42 epitopes localized in the middle and C-terminal regions defined by Aβ residues 16-30, 19-33 and 28-42 in an HLA-DR-restricted manner^4,5^. Yet, and starting from human peripheral blood mononuclear cells (PBMCs), the authors found T-cell lines reactive to the Aβ1-28 region. Although the response elicited by this specific site was less frequent -2 of 24 PBMC-derived T-cell lines-, it was HLA-DR-dependent and supported the presence of a subset of T-cells with reactivity against the Aβ(1-28) N-terminal fraction.

Additional studies by Kutzler and colleagues provided further support and functional evidence for HLA-DR1-mediated recognition of Aβ N-terminal region using HLA-II transgenic mice^6^. Immunization with human Aβ1-42 elicited IFN-γ responses in a manner dependent on Aβ peptides, and immunologically predominant within Aβ(1-28). Importantly, this reactivity was sustained regardless of CD8^+^ T-cell depletion, thus suggesting a CD4^+^ T-cell driven response to Aβ and mediated by HLA-II. In conclusion, despite human studies put the focus on the Aβ middle and C-terminal regions, the study by Kutzler *et al* support a role for Aβ(1-28) as an antigen source for HLA-DR1-display associated with CD4^+^ T-cell response.

Under these premises, an atomic view of an Aβ-derived peptide presented by HLA-DR1 contributes structural competence at the experimental level for Aβ presentation by HLA.

Aβ1-15 acquires a very specific register within the HLA-DR1 binding cleft, with Aβ residues F4 to V12 occupying the positions P1 throughout P9 that define the core, whereas residues from flanking regions extend at both ends. This arrangement is consistent with the classical open-ended binding mode observed for MHC-II molecules and highlights the contribution of peptide residues beyond the central 9-mer core. The electron density that is observed justifies the peptide binding register within the groove. The peptide is anchored to HLA-DRA and HLA-DRB by a network of polar contacts distributed along the groove. In addition, the pi-pi interaction between HLA-DRB1 Tyr59 and the peptide His at P10 suggests that the residues beyond the P1-P9 core contribute to the overall stability of the HLA-peptide assembly and shape also the surface topology targeted by a potential TCR. This is relevant because flanking residues involved in TCR recognition can determine the ultimate TCR-pMHC docking geometry.

A sequence scan for polymorphisms in HLA-DRB suggests that specific variations in alleles could influence the binding of beta-amyloid peptide. One position at the peptide-HLA binding interface, DRB70, was noticed to be polymorphic and potentially relevant. Position 70 in HLA-DRB1*01:01 is occupied by Arg, whose side chain is projected towards the peptide binding interface. Arg70 contacts Aβ1-15 through multiple interactions: a salt bridge with Asp (P4) and via H-bond with Ser (P5) backbone. This scenario supports a relevant role for DRB1 Arg70 in the overall accommodation and register of Aβ1-15 within HLA-DR1*01:01. Alignment of HLA-DRB1 alleles associated with risk or protection in AD^7^ underscores this position as a potentially relevant polymorphism. Variations at this position, observed throughout the aligned HLA-DRB alleles, are expected to have an impact in the structural stability of the HLA:peptide assembly. Substitution to Lys in DRB1*04:01could be conservative for electrostatic reasons. However, replacement by Ala in DRB1*15:01 would alter the local charge and consequently the interaction with Asp (P4), whereas Glu, as seen in DRB1*13:02, would represent a more drastic change due to provision of a fully opposite electrostatic charge. Since Arg70 is present in several other alleles reported as protective^7^, this residue represents a polymorphic contact that could dictate accommodation in HLA-DRB.

The HLA-DRB1*01:01:Aβ1-15 crystal structure does not establish per se that Aβ1-15 represents an immunodominant Aβ epitope associated with AD pathogenic T-cell reactivity. Instead, it settles an Aβ-derived fragment that can accommodate within HLA-DR1 in a structurally coherent manner. Structural competence for the binding of a particular peptide is a property that does not directly imply the natural generation of such antigen nor does it translate into T-cell reactivity to that particular peptide.

Therefore, this study does not claim Aβ1-15 as a naturally processed antigen presented by microglia-resident HLA-II. We do not provide either evidence for CD4^+^ T-cell recognition of Aβ1-15. Consequently, the HLA-DRB1*01:01:Aβ1-15 structure should not be interpreted as a functional evidence for Aβ1-15 presentation to T-cells, but as an experimentally-determined molecular framework providing an ordered and plausible pMHC topology candidate for T-cell recognition. Because of their electrostatic nature and structural accesibility, as observed in the present structure, Arg (P2), His (P3), Ser (P5) and Glu (P8) are plausible Aβ-T-cell recognition motifs. Arg (P2) and Ser (P5) side chains are involved in MHC engagement. Consequently, their availability for T-cell targeting is less likely, although conformational rearrangements leading to their involvement and enable TCR docking is an open possibility.

In summary, this study brings the first atomic view of an Aβ fragment bound to an HLA molecule and settles Aβ1-15 as an structurally coherent antigenic candidate for HLA presentation. The peptide binding register and the peptide-MHC contacts that sustain a potential TCR-binding motif are described.

## Methods

### Peptide binding prediction

Prediction of Aβ42-derived peptides to HLA alleles was carried out according to NetMHCIIpan - 4.2 parameters^8^. Human Aβ42 was used as source of 15-mer peptides. Binding parameters were assessed against HLA-DRB1*01:01, *03:01, *04:01, *13:02 and 15:01. Peptide candidates are labeled as weak or strong binders following NetMHCIIpan 4.2 eluted-ligand rank thresholds, i.e., ≤ 1% for strong binders, and ≤ 5% for weak binders.

### Protein production

The HLA-DRB1*01:01:Aβ1-15 was recombinantly produced in CHO cells. HLA-DRA and HLA-DRB1*01:01 genes were synthesized by GeneUniversal. Aβ1-15 was fused upstream HLA-DRB1*01:01 through a glycine-serine linker, and the HLA-DRA and HLA-DRB1 native chains were extended with a 3C rhinovirus site followed by knob and hole sequences, respectively, to enhance protein yields, as described previously^9^. A C-terminal Twin-Strep-tag® (IBA Lifesciences) was included in the HLA-DRA construct for downstream purification steps. The HLA heterodimer with the bound peptide was isolated from the cell culture supernatant. First, the cell supernatant was freed of cells by centrifugation, filtered and buffer exchanged to 100 mM Hepes pH 7.4, 150 mM NaCl (HBS), 1 mM EDTA. Following this, the protein was captured in a 5-ml Strep-Tactin XT 4Flow (Fisher Scientific) column, the contaminants removed with a washing step, and the target protein eluted in HBS, 50 mM biotin. The sample was left at 4 ºC overnight in the presence of in-house made 3C^10^ at a 1:100 enzyme:substrate (weight:weight) ratio for removal of knob and hole extensions. The digested sample was buffer exchanged to HBS and loaded onto a 1-ml Strep-Tactin XT 4Flow (Fisher Scientific) column. The flow-through was recovered, HLA-DRB1*01:01:Aβ1-15 further purified through a Superdex 200 10/300 GL column (Cytiva) column. The final protein product was concentrated to 20 mg/ml for crystallization purposes.

### Protein crystallization

HLA-DRB1*01:01:Aβ1-15 crystals appeared after two months at 22 ºC in 0.2 M MgCl_2_, 0.1 M Tris pH 8.5, 25% PEG 3350 using sitting drop plates with protein-reservoir mixtures of 0.25 microliters each. Crystals were cryo-cooled in the presence of 20% ethylenglycol.

### Structure solution

Full X-ray diffraction datasets were collected at Xaira beamline (Cerdanyola del Vallès, Barcelona, Spain). Diffraction data were processed with a DIALS/Xia2 package^11,12^, then scaled and merged with Aimless^13^. The structure was solved with Phaser using as template the HLA-DRB1*01:01 model generated with Alphafold^14^. Refinement was carried out with Refmac5^15^ and Phenix.refine^16^ with non-crystallographic symmetry (NCS), and the Aβ1-15 peptides were built for both HLA-DR1 molecules present in the asymmetric unit, according to positive Fo-Fc signal. The structure was further built and polished through several cycles of model building in Coot^17^ and further refinement.

## Acknowledgements

The authors are grateful to the Memorial consortium, the staff of Xaira and Xaloc beamlines at ALBA Synchrotron for their assistance with X-ray diffraction data collection and the wwPDB Biocuration Staff for their support in the deposition of atomic coordinates and structure factors in the Protein Data Bank.

## Autorship contributions

Research conception and supervision: JLS; Experimental work: EE, GDR, LO and JLS. Manuscript writing: JLS.

## Data availability

Atomic coordinates and structure factors have been deposited and are available at the Protein Data Bank under the accession code PDB ID 31VO.

